# Cell-type specific mechanical response and myosin dynamics during retinal lens development in *Drosophila*

**DOI:** 10.1101/558593

**Authors:** Laura Blackie, Rhian F. Walther, Michael F. Staddon, Shiladitya Banerjee, Franck Pichaud

## Abstract

During organogenesis, different cell types need to work together to induce functional multicellular structures. To study this process, we made use of the genetically tractable fly retina, with a focus on the mechanisms that coordinate morphogenesis between the different epithelial cell types that make up the optical lens. Our work shows that these epithelial cells present contractile apical-medial MyosinII meshworks, which control the apical area and junctional geometry of these cells during lens development. Our study also suggests that MyosinII meshworks drive cell shape changes in response to external forces, and thus they mediate part of the biomechanical coupling that takes place between these cells. Importantly, our work, including mathematical modelling of forces and material stiffness during lens development, raises the possibility that increased cell stiffness acts as a mechanism for limiting this mechanical coupling. We propose this might be required in complex tissues, where different cell types undergo concurrent morphogenesis and where averaging out of forces across cells could compromise individual cell apical geometry and thereby organ function.

## INTRODUCTION

Many of our organs consist of different polarized cell types, including epithelial cells, which adhere to one another through lateral adherens junctions (AJ) to form tissues. How different cell types work together to induce a complex tissue to generate a functional organ is not fully understood. To a large extent, epithelial tissue patterning has mostly been studied in relatively simple, homogeneous epithelia that consist of one cell type, with a focus on specific instances of cell shape change such as during apical constriction (Martin *et al.*, 2009a), cell intercalation in the fly embryo (Bertet *et al.*, 2004; Blankenship *et al.*, 2006) or cell spreading during zebrafish gastrulation (Lavoie *et al.*, 2017). In these simple epithelia, cell and tissue shape depends in part on the balance of contractile forces generated by the actomyosin cytoskeleton and intercellular adhesion through Cadherins (Heisenberg and Bellaiche, 2013; Munjal and Lecuit, 2014; Lecuit and Yap, 2015). In this balance, adhesion promotes AJ extension while Myosin-II (MyoII) contractility antagonizes this process.

An essential regulator of cell and tissue shape is the contractile actomyosin cytoskeleton, which consists of at least two pools – a medial meshwork that runs below the apical membrane, and actomyosin filaments that localize at the AJ. These two pools are linked, as the medial meshwork is anchored to the AJ through discrete points of contact (Roh-Johnson *et al.*, 2012; Heisenberg and Bellaiche, 2013). The medial meshwork is pulsatile, with cycles of discrete node contraction and relaxation that occur over tens of seconds, and that can promote apical area fluctuations and AJ remodelling over similar time scales (Coravos *et al.*, 2017). Contractile pulses appear to be self-organizing and associated with cycles of phosphorylation and dephosphorylation of the MyoII regulatory light chain, and thus cycles of assembly/disassembly of the meshworks (Kasza *et al.*, 2014; Vasquez *et al.*, 2014; Munjal *et al.*, 2015; Mason *et al.*, 2016). The AJ pool of actomyosin is linked to the Cadherin system and is thought to function as part of a ratchet mechanism that can harness the contractile forces applied onto the AJ by the medial meshwork to promote AJ remodelling (Coravos *et al.*, 2017). As morphogenesis proceeds in a simple epithelium, individual cell heterogeneities tend to be ‘averaged out’ to produce a homogenous tissue, consisting of cells with very similar apical geometry (Gibson *et al.*, 2006; Farhadifar *et al.*, 2007).

However, most organs consist of complex tissues where different cell types adopt distinct apical geometries that best suit their function. How different cell apical geometries and areas are generated within a group of cells, and what mechanisms mediate their persistence despite exposure to forces arising from concurrent cell morphogenetic programs remain poorly understood. Recent studies have begun to investigate these questions by studying the mechanical properties of different tissues undergoing concurrent morphogenesis, and how cell packing is organised in 3D. These studies have revealed key features of epithelial morphogenesis. In the fly wing for example, the actomyosin cytoskeleton can reorganise to increase tissue stiffness in response to extrinsic tensile forces (Duda *et al.*, 2019). Similarly, different mechanical properties correlate with different tissue behaviours in the fly embryo (Rauzi *et al.*, 2015). Cell shape and packing have also been shown to be constrained in a curved environment and when tissues are bent (Rupprecht *et al.*, 2017; Gomez-Galvez *et al.*, 2018). All these studies point to an important role for mechanical forces during tissue morphogenesis.

Here we made use of the *Drosophila* retina to study the mechanical properties of different cell types as they undergo distinct, concurrent morphogenesis programs to assemble into a functional multicellular unit. All retinal cells are genetically tractable and can be imaged at high spatial-temporal resolution intra-vitally (Fichelson *et al.*, 2012). The fly retina consists of approximately 750 identical physiological units called ommatidia. Each ommatidium comprises different epithelial cell types: four cone cells, surrounded by two primary pigment cells, themselves surrounded by a ring of interommatidial cells (Ready, 1989) (Figure 1A). During ommatidium morphogenesis, these epithelial cells acquire distinct apical geometries and areas. Similar to simple epithelia, the apical geometry of retinal cells is regulated by a combination of adhesion and actomyosin contractility at the AJ (Del Signore *et al.*, 2018). In particular, the interplay between E/Ncadherin expression in the cone cells controls their apical geometry and topology in the plane of the epithelium (Hayashi and Carthew, 2004; Kafer *et al.*, 2007; Chan *et al.*, 2017). This function for NCadherin has been linked to the downregulation of MyoII accumulation at the AJ shared between the cone cells and increased MyoII levels at the AJ these cells share with the surrounding pigment cells (Chan *et al.*, 2017). While adhesion and actomyosin at the AJ work together to pattern the optical lens, the medial/ cytoplasmic actomyosin cytoskeleton of the corresponding retinal cells has not been investigated in detail. In addition, it is not clear how forces are balanced across the ommatidium as the lens cells undergo morphogenesis. Here we combined molecular genetics, light-induced perturbation experiments and computational modelling to describe the MyoII cytoskeleton in retinal epithelial cells that make up the optical lens, and to probe the balance of forces that are at play during eye development. Our work shows that all lens cell types present medial MyoII meshworks. These are contractile but do not show any particular persistent polarization in the way they contract. Our results indicate they contribute to controlling the shape and size of the apical area of retinal cells. They also drive cell shape changes in response to force perturbation in all lens cell types but one: the lens secreting cone cells, which we find keep their apical area and shape largely unchanged when challenged by extrinsic forces. Generation of a 2D mechanical model of forces in the developing lens and genetic perturbation of the contractile actomyosin cytoskeleton suggest that this is because cone cells are stiffer than the other retinal cell types. Our work thus reveals that cell stiffness is an important parameter of tissue morphogenesis that modulates the ability of cells to respond to mechanical forces.

**Figure 1:**
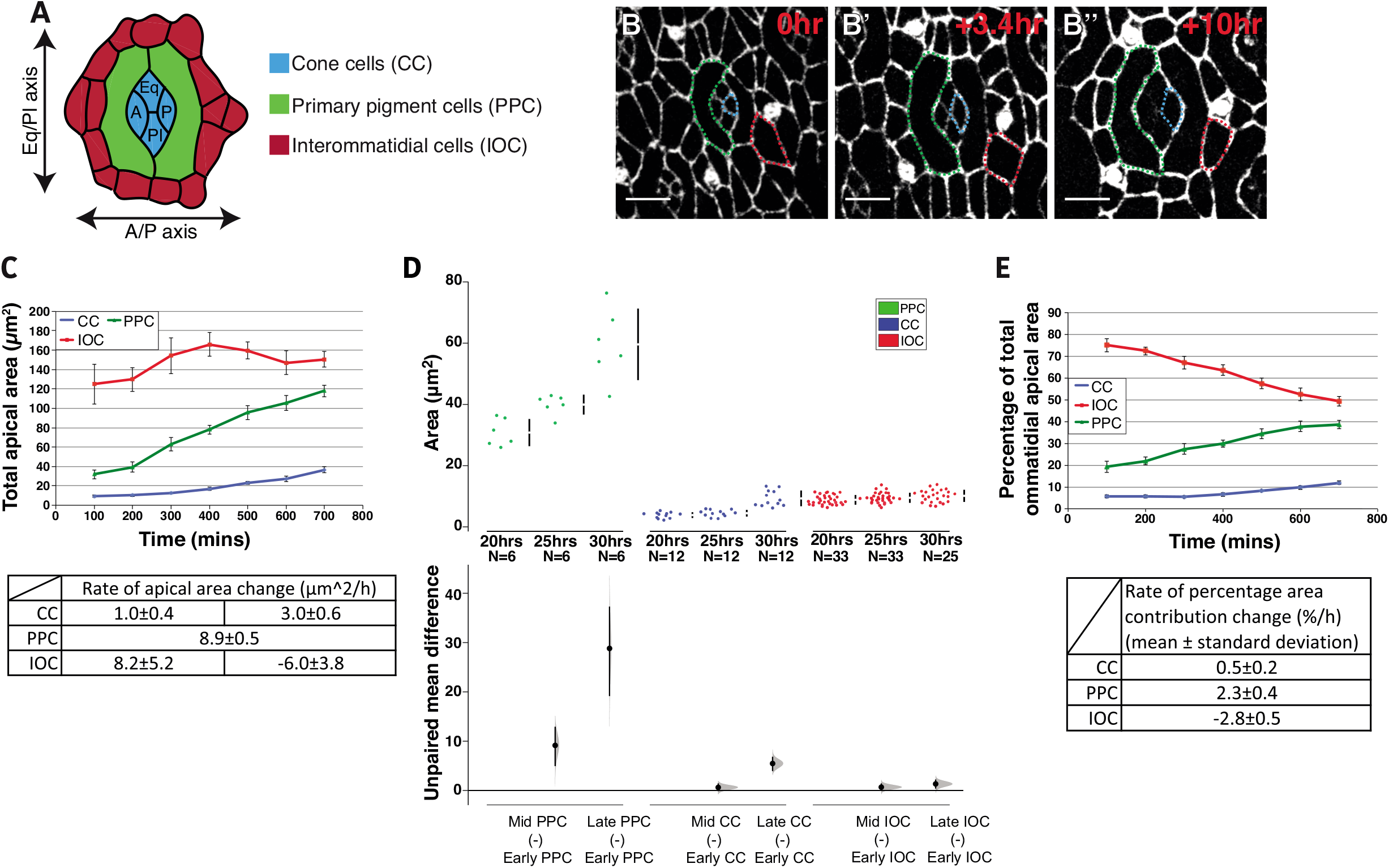
Retinal cells expand their apical area during lens development. **(A)** The arrangement of cells in the ommatidium. A = anterior, P = posterior, Eq = equatorial and Pl = Polar. **(B-B’’)** Snapshots taken from a time-lapse movie of ommatidium development, with AJs labelled with endogenous Ecad::GFP. Scale bar = 5μm. **(C)** Total apical area of each of the three cell layers over time (n=13 ommatidia from 2 pupae). Table indicates rates of apical area change over time. Error bars = standard deviation (S.D.) **(D)** Cummings estimation plot with upper axis showing distribution of apical area of individual primary pigment cells, cone cells and interommatidial cells at three different stages of morphogenesis (n=9 ommatidia from 5 pupae). On the lower axis, mean differences for comparisons to the cell apical area at 20hrs are plotted as bootstrap sampling distributions, dot=mean difference, error bars = 95% confidence interval. Unpaired mean difference of: MidPPC (n=6) minus EarlyPPC (n=6): 9.14 [95CI 4.94; 12.9]; LatePPC (n=6) minus EarlyPPC (n=6): 28.8 [95CI 19.1; 37.3]; MidCC (n=12) minus EarlyCC (n=12): 0.616 [95CI −0.141; 1.43]; LateCC (n=12) minus EarlyCC (n=12): 5.47 [95CI 3.93; 6.84]; MidIOC (n=33) minus EarlyIOC (n=33): 0.667 [95CI −0.0907; 1.42]; LateIOC (n=33) minus EarlyIOC (n=25): 1.31 [95CI 0.38; 2.22]. **(E)** Percentage apical area of each of the three cell layers relative to the apical area of the whole ommatidium over time (n=13 ommatidia from 2 pupae). Table indicates rates of percentage apical area change over time. Error bars = S.D.

## RESULTS

### Retinal cells increase their apical area during ommatidium morphogenesis

Firstly, we sought to examine the dynamics of apical area changes in retinal cells that make up the lens as they acquire their position and apical geometry during ommatidium development (Figure 1B, Movie S1). During ommatidium morphogenesis, the total apical area of the ommatidial core cells, defined as the cone and primary pigment cells, increases over time (Larson *et al.*, 2010). We examined 13 ommatidia using time-lapse imaging, ensuring they were of similar developmental stage. These quantifications confirmed a previous report that the total apical area of the core cells increases through time (Figure 1C) (Larson *et al.*, 2010). It also showed that the combined apical area of the interommatidial cells increases transiently, then decreases as these cells thin and some are eliminated through apoptosis (Brachmann and Cagan, 2003) (Figure 1C). Additionally, we found that as retinal cells gradually remodelled their apical area and shape, they did so at different, cell type specific rates (Figure 1C). The primary pigment cells had an average rate of area change of 8.9 ± 0.5 μm^2^/h (mean ± standard deviation (S.D.)). The cone cells and interommatidial cells had two phases of apical area change. For the cone cells, we found that there was an initial slow expansion phase of 1.0 ± 0.4 μm^2^/h, followed by a faster phase of 3.0 ± 0.6μm^2^/h.

Interommatidial cells had an initial fast expansion rate of 8.2 ± 5.2 μm^2^/h on average, followed by a slower average rate of area decrease of −6.0 ± 3.8 μm^2^/h. Thus, the rate of the initial phase of area expansion in the interommatidial cells is very similar to that of the primary pigment cells. Examining the changes in area of individual cells over time (Figure 1D) showed that they follow a similar trend to that of the respective cell type groups. Consistent with these results, analysis of the relative contributions of the cone cells, primary pigment cells and interommatidial cells to total ommatidial lens area demonstrated that the contribution of the cone and primary pigment cells increases over time while that of the interommatidial cells decreased (Figure 1E, Movie S1).

### Apical area fluctuations correlate with apical expansion rates

In simple epithelia, changes in apical area have been shown to occur through small fluctuations that occur over short time-scales and are stabilized incrementally (Martin *et al.*, 2009a; Fernandez-Gonzalez and Zallen, 2011; Sawyer *et al.*, 2011). Here, we tested whether this is the case in the retina. We found that in the lens, all cell types undergo fluctuations in apical area over a few minutes (Figure 2A). However, the amplitude of area fluctuations was different for each cell type (Figure 2B). The primary pigment cells exhibited the largest fluctuations followed by the interommatidial and cone cells (Figure 2B-C). As cell surface stiffness is inversely proportional to shape fluctuations (Turlier, 2019), our results suggest that the cone cells are the stiffest ommatidial cell type. Normalising the area of individual cells by their average revealed that these differences in fluctuation between cell types are linked to their area (Figure 2D). Thus, the primary pigment cells, which have the largest apical area also have the highest median area fluctuation amplitudes. To complement this analysis, we also quantified the cycle length of fluctuations (defined as the time between subsequent peaks). We found that these cycle lengths were similar between retinal cell types (Figure 2E). Altogether, these quantifications suggest the cells that have the largest apical area and highest fluctuations over short time scales, are the cells that undergo the fastest expansion of their apical area through long time scales.

**Figure 2:**
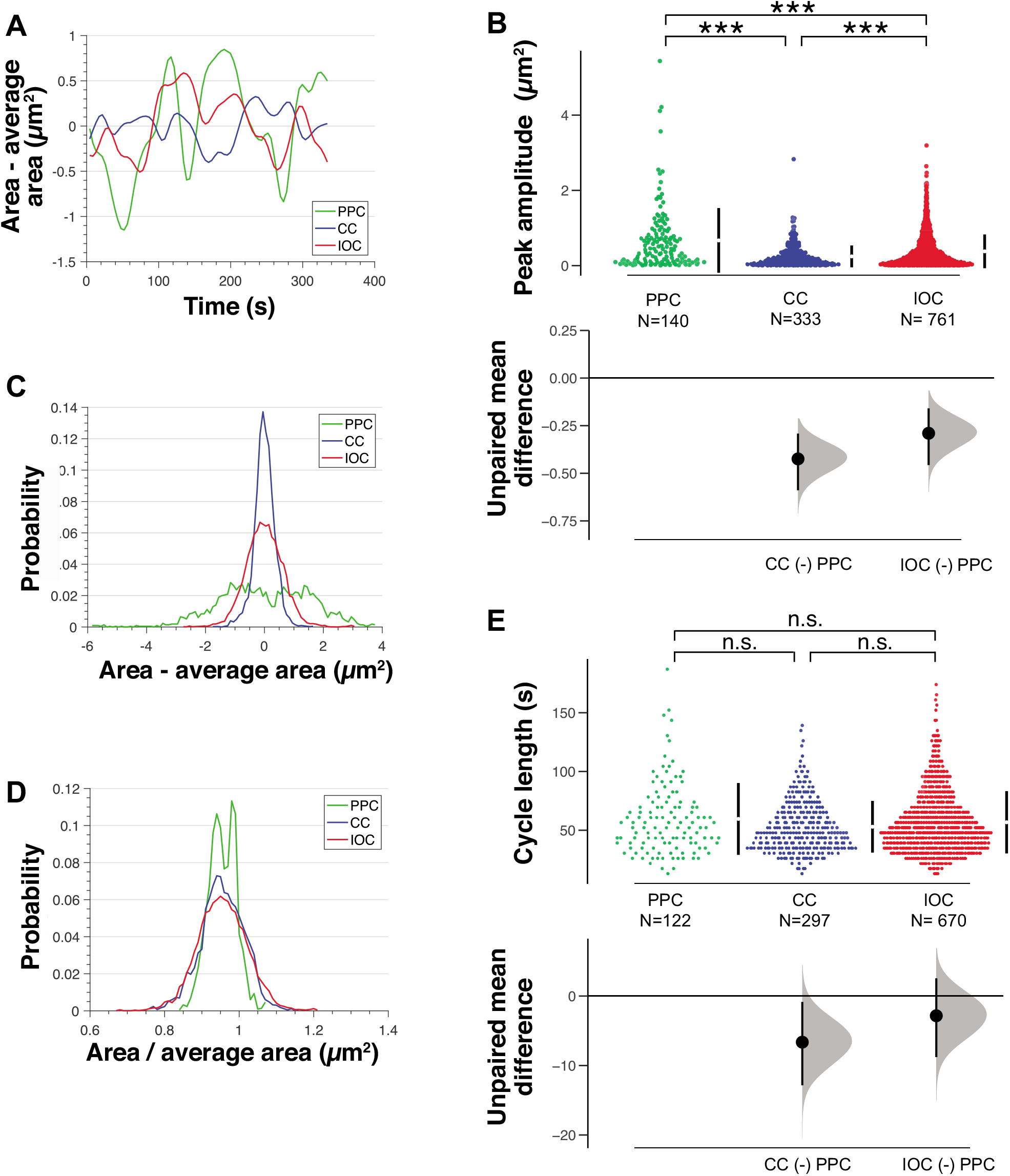
Highest apical area fluctuations correlate with fastest apical expansion rates. **(A)** Apical area minus average apical area for each cell type over time for one representative primary pigment, cone and interommatidial cell. **(B)** Cummings estimation plot with upper axis showing distribution of amplitude of peaks in area fluctuations for primary pigment, cone and interommatidial cells (n=9 ommatidia from 5 pupae, Kruskal-Wallis test, p<0.0001, post-hoc Dunn’s multiple comparisons tests: primary pigment-cone cells p<0.0001, primary pigment-primary pigment p<0.0001, cone-primary pigment p=0.0004). On the lower axis, mean differences for comparisons to PPC are plotted as bootstrap sampling distributions, dot=mean difference, error bars = 95% confidence interval. Unpaired mean difference of: CC (n=140) minus PPC (n=333): −0.424 [95CI −0.589; −0.291]; IOC (n=140) minus PPC (n=761): −0.29 [95CI – 0.457; −0.159] **(C)** Probability distribution of apical area minus average apical area for primary pigment, cone and interommatidial cells (n=9 ommatidia). **(D)** Probability distribution of apical area normalized by average apical area for primary pigment, cone and interommatidial cells (n=9 ommatidia). **(E)** Cummings estimation plot with upper axis showing distribution of cycle lengths of area fluctuations for primary pigment, cone and interommatidial cells (n=9 ommatidia from 5 pupae, Kruskal-Wallis test, p=0.1708). On the lower axis, mean differences for comparisons to PPC are plotted as bootstrap sampling distributions, dot=mean difference, error bars = 95% confidence interval. Unpaired mean difference of: CC (n=122) minus PPC (n=297): −6.64 [95CI – 12.8; −0.846]; IOC (n=122) minus PPC (n=670): −2.84 [95CI −8.78; 2.55].

### Myosin pulse contraction correlates with fluctuations in cell area

During cell apical constriction, the medial actomyosin meshworks associated with the apical surface of the cell drive the cell’s area fluctuation (Martin *et al.*, 2009b; Fernandez-Gonzalez and Zallen, 2011; Sawyer *et al.*, 2011). In order to determine whether this is also the case when considering fluctuations of retinal cell area, we examined the localization and dynamics of MyoII using a GFP-tagged version of the regulatory light chain, Sqh::GFP (Royou *et al.*, 2002). We found that Sqh::GFP localized to the cell contacts and to a medial meshwork in all retinal lens cells (Figure 3A, Movie S2). Multiple nodes of high Sqh::GFP intensity could arise simultaneously at different locations in the cytosol of retinal cells (Figure 3B). Further, the percentage of cell area occupied by MyoII nodes and the number of nodes was highest in the interommatidial cells, which undergo apical area thinning, compared to the primary pigment or cone cells (Figure 3C-C’).

**Figure 3:**
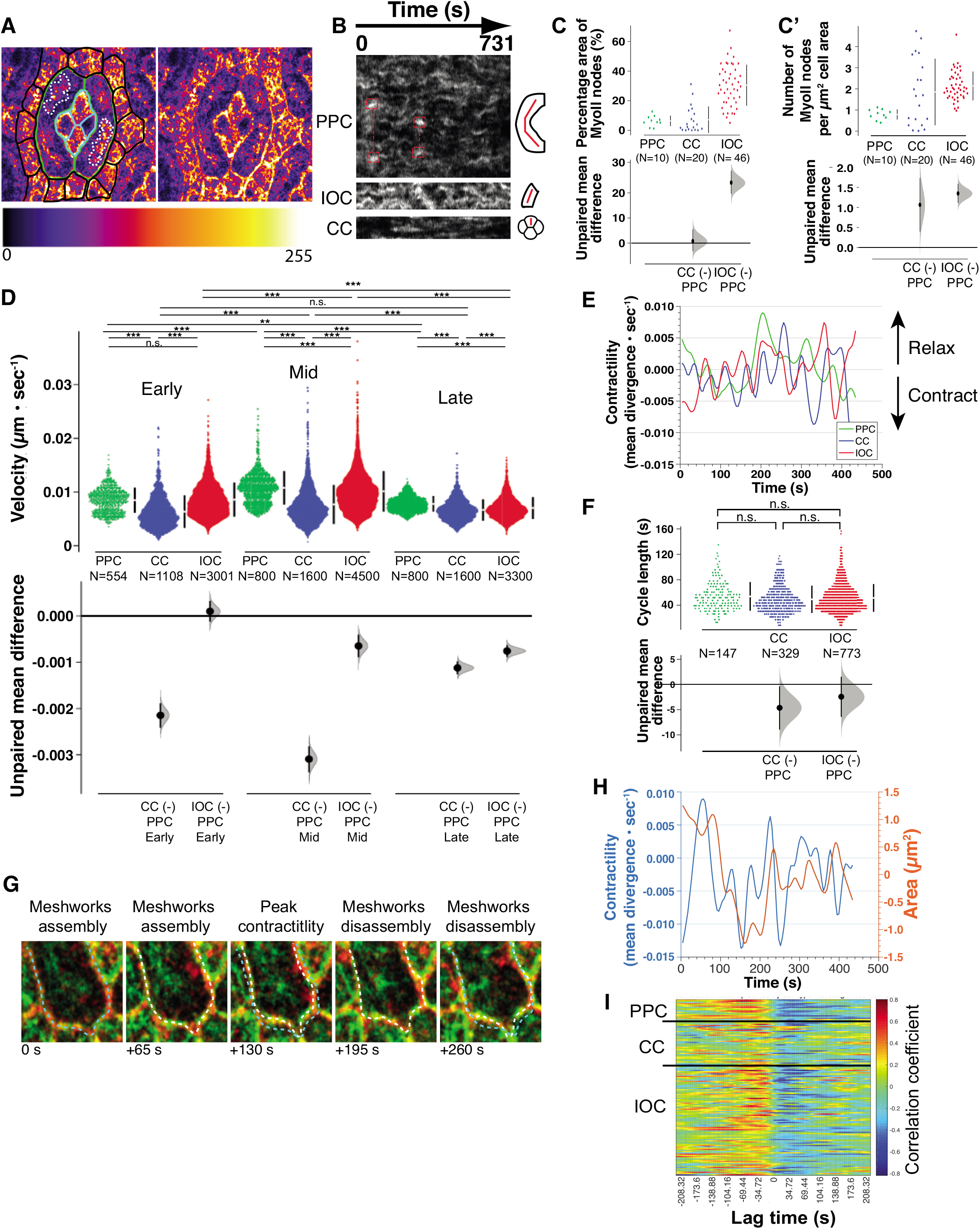
Myosin pulse contraction correlates with cell area fluctuation. **(A)** *sqh^AX3^; sqhGFP;* ommatidium showing that MyoII localizes to an extensive medial meshwork across all cell types. Interommatidial cells are outlined in black, the primary pigment cells in green and cone cells in turquoise. High-intensity MyoII meshworks are outlined by a dashed line in the primary pigment cells. Intensity of MyoII ranges from low (black) to high (white). **(B)** Kymographs showing MyoII intensity along a line through the center of each cell, indicated by red line in diagram, for each cell type over time along the X-axis. Red outlines show examples of two nodes occurring concurrently in the same cell. **(C-C’)** Density of high intensity MyoII nodes measured by **(C)** percentage of cell area covered by MyoII nodes and by **(C’)** count of number of MyoII nodes per μm^2^ of cell area. **(C)** Cummings estimation plot with upper axis showing distribution of percentages of cell area covered by MyoII nodes. On the lower axis, mean differences for comparisons to PPC are plotted as bootstrap sampling distributions, dot=mean difference, error bars = 95% confidence interval. Unpaired mean difference of: CC (n=10) minus PPC (n=20): 0.771 [95CI −3.2; 5.48]; IOC (n=10) minus PPC (n=46): 24.1 [95CI 19.7; 28.8]. **(C’)** Cummings estimation plot with upper axis showing distribution of counts of MyoII nodes per cell area. On the lower axis, mean differences for comparisons to PPC are plotted as bootstrap sampling distributions, dot=mean difference, error bars = 95% confidence interval. Unpaired mean difference of: CC (n=10) minus PPC (n=20): 1.07 [95CI 0.397; 1.74]; IOC (n=10) minus PPC (n=46): 1.35 [95CI 1.12; 1.59]. **(D)** Cummings estimation plot with upper axis showing distribution of velocities of MyoII medial meshwork at each stage of morphogenesis for primary pigment, cone and interommatidial cells (n=9 ommatidia from 5 pupae, Kruskal-Wallis and Dunn’s post-hoc tests, p<0.0001 for all tests except primary pigment-primary pigment at 20hrs APF where p=0.5545). On the lower axis, mean differences for comparisons to PPC at each stage of morphogenesis are plotted as bootstrap sampling distributions, dot=mean difference, error bars = 95% confidence interval. Unpaired mean difference of: CCearly (n=554) minus PPCearly (n=1108): – 0.00215 [95CI −0.00241; −0.00189]; IOCearly (n=554) minus PPCearly (n=3001): 9.96e-05 [95CI −0.000123; 0.000323]; CCmid (n=800) minus PPCmid (n=1600): – 0.00309 [95CI −0.00337; −0.00282]; IOCmid (n=800) minus PPCmid (n=4500): – 0.000648 [95CI −0.00089; −0.000407]; CClate (n=800) minus PPClate (n=1600): – 0.00112 [95CI −0.00126; −0.000984]; IOClate (n=800) minus PPClate (n=3300): – 0.000758 [95CI −0.000882; −0.000636], using 10000 bootstrap resamples. **(E)** Contractility (mean divergence) over time for one representative primary pigment, cone and interommatidial cell. **(F)** Cummings estimation plot with upper axis showing distribution of cycle lengths of peaks in contractility fluctuations for primary pigment, cone and interommatidial cells (n=9 ommatidia from 5 pupae, Kruskal-Wallis test, p=0.053). On the lower axis, mean differences for comparisons to primary pigment cell are plotted as bootstrap sampling distributions, dot=mean difference, error bars = 95% confidence interval. Unpaired mean difference of: CC (n=147) minus PPC (n=329): – 4.63 [95CI −8.95; −0.36]; IOC (n=147) minus PPC (n=773): −2.43 [95CI −6.42; 1.54]. **(G)** High magnification images of a pulse of MyoII assembling and disassembling in a primary pigment cell. MyoII is labelled with Sqh::GFP (green) and *AJs* are labelled with Ecad::Tomato (red). The AJ of the primary pigment cell is outlined with a dashed line. A turquoise line outlines the AJ at the onset of contraction and is used for reference in the subsequent panels. A white dashed line is then used to outline the AJ as the cell undergo local pulsed-contraction. Note how primary pigment cell contracts as a MyoII pulse assembles and then relaxes as the pulse disassembles. **(H)** Fluctuations of MyoII contractility and apical area for one representative interommatidial cell. Note that the peak in MyoII contractility precedes the peak of apical area. Frame interval in the corresponding time lapse was 4.34 sec. **(I)** Heatmap showing temporal crosscorrelation for multiple primary pigment, cone and interommatidial cells. This analysis was performed using calculated mean divergence for each cell type, averaged across the apical area of the cell. Each row represents an individual cell (n=9 ommatidia).

Next, to examine the dynamics of the MyoII meshworks in each cell type, we used Particle Image Velocimetry (PIV) (Tseng *et al.*, 2012). First, we calculated the average advection speed of Sqh::GFP particles within the medial meshwork, excluding the pool of Sqh::GFP at the AJ. We found that the velocity of Sqh::GFP motion varies between cell types and between developmental stages (Figure 3D). At early stages, Sqh::GFP particles had similar median velocity in the primary and interommatidial cells, which was greater than that estimated in the cone cells. At mid and late stages, the primary pigment cells presented the highest median velocity followed by the interommatidial and then the cone cells. Therefore, as lens development proceeds, cells with highest average area fluctuation amplitudes over short time scales (Figure 2B) also present the highest median velocity of Sqh::GFP particle displacement. In addition, the median velocity of Sqh::GFP particle displacement increased for all cells up to when they have acquired their mature geometry (20-25hrs after pupal formation (APF)), at which point they slowed down (at 30hrs APF) (Figure 3D).

We then used these PIV analyses to estimate the contractility of the actomyosin networks. This was done by calculating the divergence of the corresponding vector fields. Retinal cells present large distributed meshworks (Figure 3A and Movie S2). In order to extract trends at the cell level, we spatially averaged the calculated divergences for each cell type, excluding the junctional MyoII. This approach revealed that the estimated contractility of the cell medial meshworks fluctuates in all cell types Q with cyclical periods of contraction (pulse assembly: mean divergence < 0) and relaxation (pulse disassembly: mean divergence > 0) (Figure 3E). We found that median cycle length of fluctuations in MyoII contractility was in the range of that calculated for the area fluctuation observed on short time scales (Figure 2E and Figure 3F). This observation is in good agreement with the notion that pulse contraction of the medial myosin meshwork can drive apical area fluctuations over short time scales. Further analysis comparing time spent contracting *versus* relaxing showed that on these short timescales, there was no net contraction or relaxation (Supplementary Figure 1A-B).

To further assess whether MyoII pulse contraction is linked to cell area fluctuation, we analyzed ommatidia expressing Sqh::GFP and Ecad::Tomato to label the AJ perimeters. Focusing on the primary pigment cells revealed that localized contraction of the MyoII meshwork occurred in tandem with changes in cell apical area (Figure 3G, Movie S3). Further, apical area fluctuation and MyoII contractility fluctuation were strongly cross-correlated for the primary pigment cells and interommatidial cells (Figure 3H-I). Each peak of MyoII contractility preceded a peak in apical area fluctuation (Figure 3H). The strongest correlation occurred at an average time-lag of – 31.0 ± 9.7 s (mean ± SEM) for the primary pigment cells and −28.4 ± 3.3 s for the interommatidial cells (Figure 3I). The correlation between peaks of MyoII contractility and apical area fluctuations was not as strong for the cone cells. Altogether, these results are consistent with the hypothesis that the contractile medial MyoII meshwork induces area fluctuation over short time scales in retinal cells, and that on the longer term, these fluctuations drive apical area changes to shape cells during lens formation.

### Medial MyoII meshworks control the apical geometry of retinal cells

Next, to more directly assess the role of MyoII contractility in controlling the apical geometry and area of the lens cells, we turned to laser ablation. We focused our work on the medial MyoII meshworks in the primary pigment cells because these cells are large and readily accessible (Figure 4A). First, we set up an ablation protocol where we could trigger a destabilization of the entire medial MyoII meshwork, without affecting the AJ pool or cell viability (Figure 4B and Movie S4-5). In these conditions, we observed a change in apical geometry and a significant increase in the apical area of the targeted cell (Figure 4C-D, Supplementary Figure 2A and Movie S4-6). To ensure that our light-induced perturbations were specific, and that the targeted cells were not irreparably damaged, we only considered experiments where the medial meshwork repaired and was re-established. Re-establishment of the medial meshwork was accompanied by a recovery in apical geometry and area of the ablated cell (Figure 4B, Movie S4).

**Figure 4:**
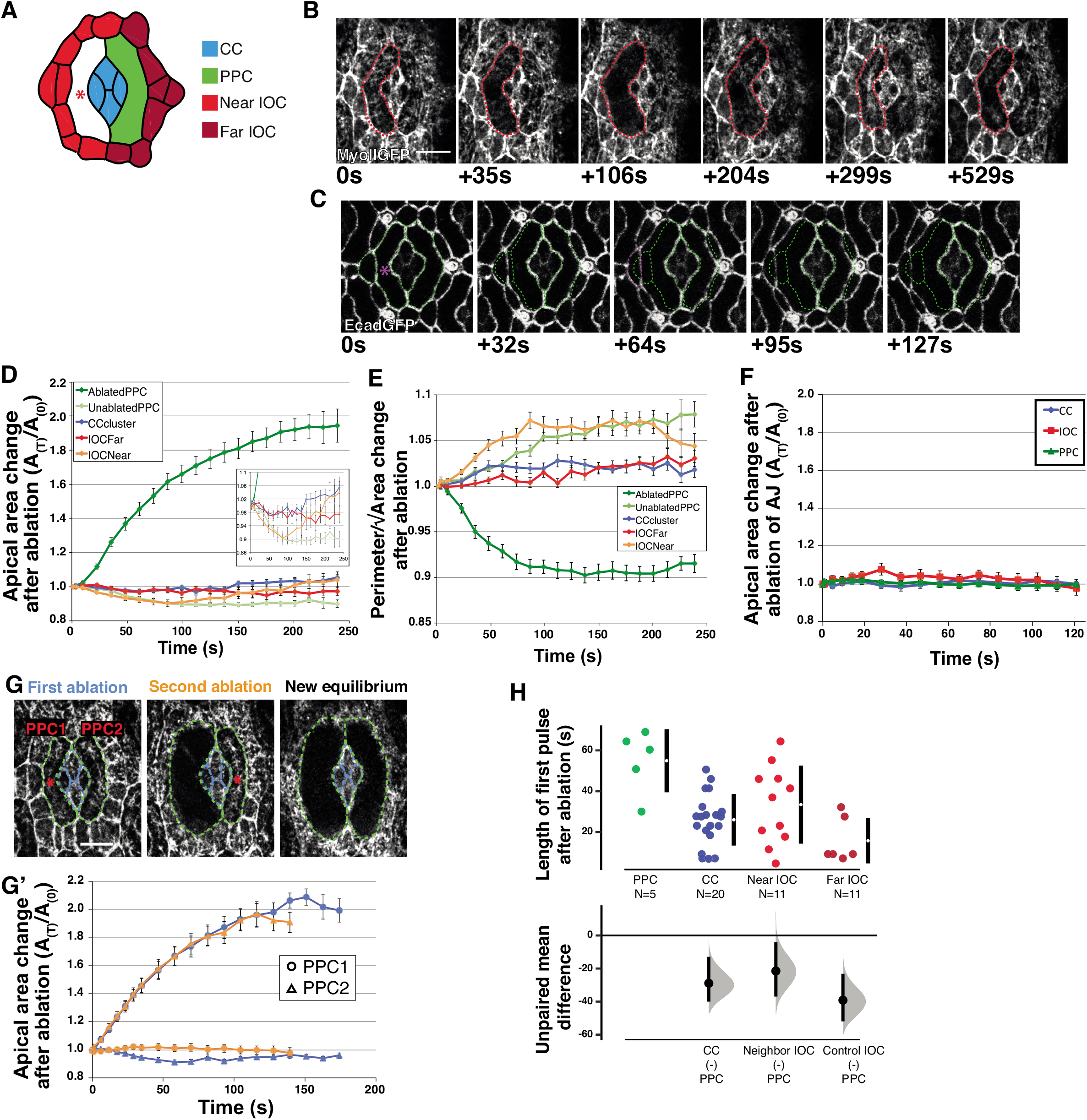
The contractile medial actomyosin meshwork controls cell shape. **(A)** Diagram of position of quantified cells relative to medial meshwork ablation (red asterisk). **(B)** Time-course showing response of cells after medial meshwork ablation in left primary pigment cell. MyoII is labelled by Sqh::GFP. Red star indicates point of laser ablation; 6 Pixels is 245nm. Ablated primary pigment cell is outlined in red. Note how cell area increases as medial meshwork is disrupted and then area is recovered as the meshwork re-establishes. **(C)** Time-course showing AJs labelled with Ecad::GFP after ablation of the medial meshwork in left primary pigment cell. Region of ablation marked with magenta star. Cell outlines before ablation superimposed on each image with green dotted line. Interommatidial cell outlined in magenta. **(D)** Change in apical area over time after ablation, A_(T)_/A_(0)_, A=apical area (n=14 ommatidia). Inset panel shows y axis at higher resolution. **(E)** Cell shape index (perimeter/√area) relative to initial shape parameter over time after ablation (n=14 ommatidia). **(F)** Apical area changes over time after ablation of the shared primary pigment cell AJ, A_(T)_/A_(0)_, A=apical area (n=5 ommatidia from 5 pupae). **(G)** Changes in apical area for two sequential ablations. The first ablation destabilizes the medial meshwork in primary pigment cell 1 (PPC1, shown in blue on the graph, **(G’)**). The second ablation destabilizes the medial meshwork in primary pigment cell 2 (PPC2, shown in orange on the graph **(G’)**). **(H)** Cummings estimation plot with upper axis showing distribution of lengths of the first MyoII contraction pulse after ablation (n=5 ommatidia from 4 pupae, one-way ANOVA p=0.0005, primary pigment cell compared to cone neighbouring interommatidial cell and control interommatidial cell: p<0.05; other comparisons: n.s.). On the lower axis, mean differences for comparisons to primary pigment cell are plotted as bootstrap sampling distributions, dot=mean difference, error bars = 95% confidence interval. Unpaired mean difference of: CC (n=5) minus PPC (n=20): −28.9 [95CI −40; −12.9]; Neighbour IOC (n=5) minus PPC (n=11): −21.5 [95CI −37; −4.06]; Control IOC (n=5) minus PPC (n=6): −39.2 [95CI −51.9; −23.2]. Error bars: D,E,F) = S.E.M. All scale bars = 5μm

### Lens cells mount cell-type specific responses to mechanical perturbation

To investigate mechanical coupling between retinal cells in these experiments, we next quantified the apical deformation of retinal cells. To this end, we measured apical area, and quantified apical geometry by calculating the shape index 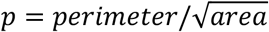, which indicates the degree of shape anisotropy (Bi, 2015). For a regular hexagon, *p* = 3.72, and increases in magnitude as the shape becomes more elongated. Ablation led to an increase in the area of the targeted cell (Figure 4B-D), and also affected the shape of its apical area (Figure 4E). In addition, the apical area of the adjacent interommatidial cells and that of the non-targeted primary pigment cell, were affected (Figure 4B-D). The area of the non-targeted primary pigment decreased, and this cell adopted a more elongated apical geometry (Figure 4D,E). This deformation, which amounted to a 10% decrease in apical area (Figure 4D), was never observed in control animals when monitoring the naturally occurring fluctuation of apical area in these cells (Supplementary Figure 2B). It is specific to the medial meshwork because ablating the AJ shared by the two primary pigment cells, did not lead to significant deformation in primary pigment cell area (Figure 4F, Movie S7). Thus, medial MyoII meshworks have a greater influence on the control of apical retinal cell area than individual junctions. Interestingly, the neighbouring cone cells did not show any change in apical area or geometry even though they share extensive AJs with the targeted cell (Figure 4D,E). Altogether, these results show that different cell types respond differently to mechanical perturbation.

**Figure 5:**
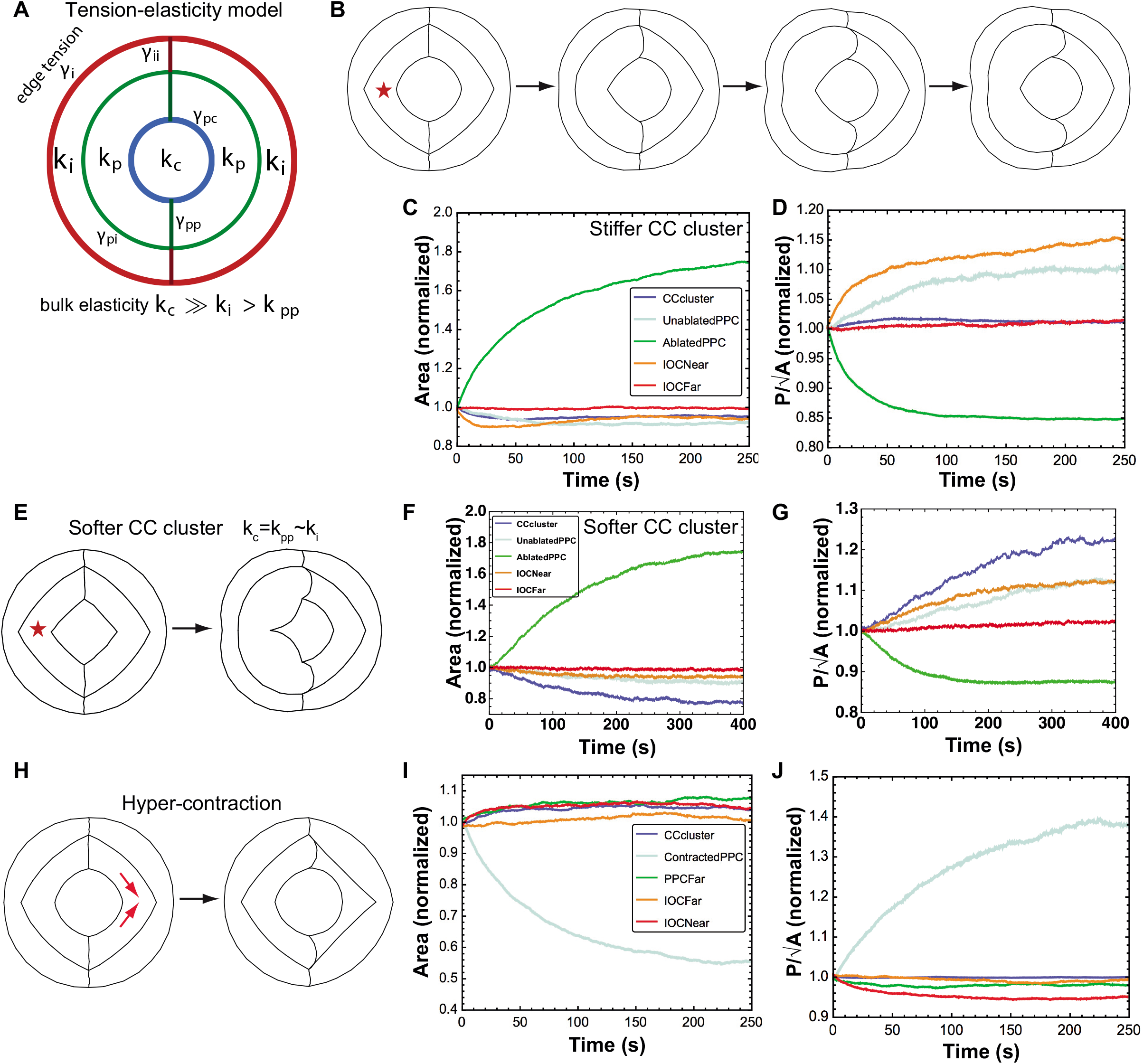
Computational model of the ommatidium predicts cell type specific mechanical response. **(A)** Schematic of the tension-elasticity model of the ommatidial cluster, showing the edge tensions, bulk elasticity parameters in the cone cell cluster (blue), primary pigment cell (green) and the interommatidial cell clusters (red). **(B)** Simulations of ablation experiments by reducing bulk tension in one of the primary pigment cells (marked by an asterisk). Left to right: evolution of colony morphology upon ablation. **(C)** Model prediction for the dynamics of the apical area (normalized) for the different cell clusters in the ommatidia, upon ablation at *t* = 0. **(D)** Model prediction for the dynamics of the cell shape index (perimeter/√area, normalized) for the different cell clusters in the ommatidia, upon ablation at *t* = 0. **(E)** Simulation of ablation experiments for a softer cone cell (CC) cluster. **(F-G)** Dynamics of cell area **(F)** and cell shape index **(G)** in the ablation simulation, showing significant deformation of the cone cell cluster, inconsistent with experimental data. **(H)** Simulation of the hypercontraction in the right primary pigment cell (marked by red arrows). **(I)** Model prediction for the dynamics of the apical area (normalized) for the different cell clusters in the ommatidia, upon the induction of hyper-contraction in the right primary pigment cell at *t* = 0. **(J)** Model prediction for the dynamics of the cell shape index (perimeter/√area, normalized) for the different cell clusters in the ommatidia, upon the induction of hyper-contraction in the right primary pigment cell at *t* = 0.

Shrinkage of the non-targeted primary pigment cell could be active via contractile forces intrinsic to the cell, or passive as a response to the expansion in apical area of the laser targeted cell. To decipher between these two possibilities, we destabilized the medial MyoII meshworks in both primary pigment cells sequentially. In this case, no decrease in apical area was observed after the second ablation (Figure 4G-G’), and both targeted cells increased their area. This confirms that apical MyoII meshworks are required to define the apical area and geometry of pigment cells. It also shows that primary pigment cell contraction in response to laser ablation requires an intact medial MyoII meshwork. Further supporting this idea, quantification of how long the first fluctuation of MyoII contraction after ablation lasts in the non-targeted primary pigment and and the near interommatidial cell revealed an increase in duration when compared to wild type ommatidia (Compare Figure 3F and Figure 4H). The primary pigment cell, which deforms the most in response to ablation, presented the highest average length of the first fluctuation of MyoII contraction after ablation (Figure 4H). Altogether, these experiments indicate that while all retinal cells are mechanically coupled though their AJ, the cone cells deform less than the other retinal cell types in response to external perturbation. Amongst the lens cells, the primary pigment cells are the most deformable in response to mechanical perturbation.

### Computational model of the ommatidium predicts that differences in cell mechanical properties coordinate cell type specific shape dynamics

To better understand the physical origin of the cell type specific mechanical response to actomyosin perturbations, we developed a two-dimensional tension-elasticity model of the ommatidium. The model construction is inspired by vertex models for epithelial morphogenesis (Fletcher *et al.*, 2014), where each cell is treated as a mechanical medium, carrying edge tensions at the cell-cell interfaces, a bulk tension arising from a contractile medial actomyosin meshwork, and area elasticity penalizing changes in the cell apical area (Methods, Figure 5A). Competition between tension and elasticity gives rise to specific cell shapes, determined by parameters characterizing edge tension, bulk tension, and area elastic modulus (Figure 5A). Since we are interested in the overall morphology of the ommatidium and its relationship to cell type specific properties, we treated the cone cell quartet as one mechanical object, each primary pigment cell as separate entities, and the left and right interommatidial cells as individual mechanical objects. Starting with an initial circular configuration for the ommatidial cluster, we dynamically evolved the points on the cell contours based on their resultant forces, in order to obtain the steady-state morphology of the ommatidium (Figure 5B). The forces evolving ommatidium morphology result from minimizing the mechanical energy of the cluster (see Methods). We benchmarked the model parameters (tension parameters, elastic moduli) to recapitulate the shapes and sizes of the control cone, interommatidial and primary pigment cells measured experimentally (Figure 1D).

As our model accurately captured ommatidial cell morphologies, we then sought to test if it could capture the results of the laser ablation experiments (Figure 4). To simulate medial meshwork ablation in the primary pigment cells, we reduced the bulk tension in the left primary pigment cell and allowed the cell cluster to dynamically evolve to a new morphology (Figure 5B). Our model quantitatively captured the experimentally measured morphological transformations, provided that the elastic modulus of the cone cell cluster (*k_c_*) is much higher than that of the interommatidial cell cluster (*k_i_*) and the primary pigment cells (*k_pp_*), i.e. *k_c_* ≫ *k_i_* > *k_pp_*. Upon reduction in bulk contractility, the area of the targeted primary pigment cell expanded (Figure 5C) while decreasing its shape index (lower aspect ratio) (Figure 5D). In agreement with experimental data (Figure 4D-E), the shape and area of the cone cell cluster remained unchanged. The interommatidial cell cluster adjacent to the targeted primary pigment cell initially decreased in area, but then recovered over time (Figure 5C), and also thinned in shape (Figure 5D). The inclusion of a rigid cone cell cluster enabled mechanical communication between the primary pigment cells, such that the untargeted primary pigment cell shrunk in area while increasing its shape index (higher aspect ratio) (Figure 5C-D). These results agree with our experiments (Figure 4D-E) and suggest that different retinal cell types have different mechanical properties. Simulations with a softer cone cell cluster, whose elasticity was comparable to primary pigment and interommatidial cells, led to significant deformations of the cone cells during ablation (Figure 5E-G), which was not observed in our experimental perturbations (Figure 4D-E). Therefore, these simulations suggest that the cone cells are stiffer than their neighbours, and that this stiffness limits their mechanical coupling.

To further test the predictive power of the model, firstly, we studied how cell shape might respond to an increase in contractility (higher bulk tension) in the right primary pigment cell (Figure 5H). One strong prediction from the model is that increasing the tension in one primary pigment cell will cause the paired primary pigment cell to increase its apical area (Figure 5I). Furthermore, our model predicted that the shape index should decrease for the paired primary pigment cell, and the interommatidial cells, while increasing for the constricted primary pigment cell (Figure 5J).

### Experimental testing of computational model predictions

To experimentally test the predictions arising from our model, we used genetics to perturb the actomyosin cytoskeleton. Firstly, to induce hypercontraction in one primary pigment cell (simulation in Figure 5H) we inhibited the expression of *sds22* (Figure 6A), which encodes for the regulatory subunit of MyoII phosphatase PP1A (Grusche *et al.*, 2009). This did not have any significant effect of the Sqh::GFP intensity within the AJs of the cell when compared to wild type suggesting this perturbation is relatively specific to the medial meshwork (Supplementary Figure 3). PIV analysis performed on Sqh::GFP demonstrated that this led to a significant decrease in medial meshwork velocity (Figure 6B-B’). This genetic perturbation resulted in a reduction in apical area and a change in geometry, whereby the targeted cell became more elongated (Figure 6A, C-D). As predicted by our computational model, this was accompanied by an increase in average apical area of the non-targeted primary pigment cell (Figure 6C). It also led to a slight decrease in the shape index for the paired cell (Figure 6D).

**Figure 6:**
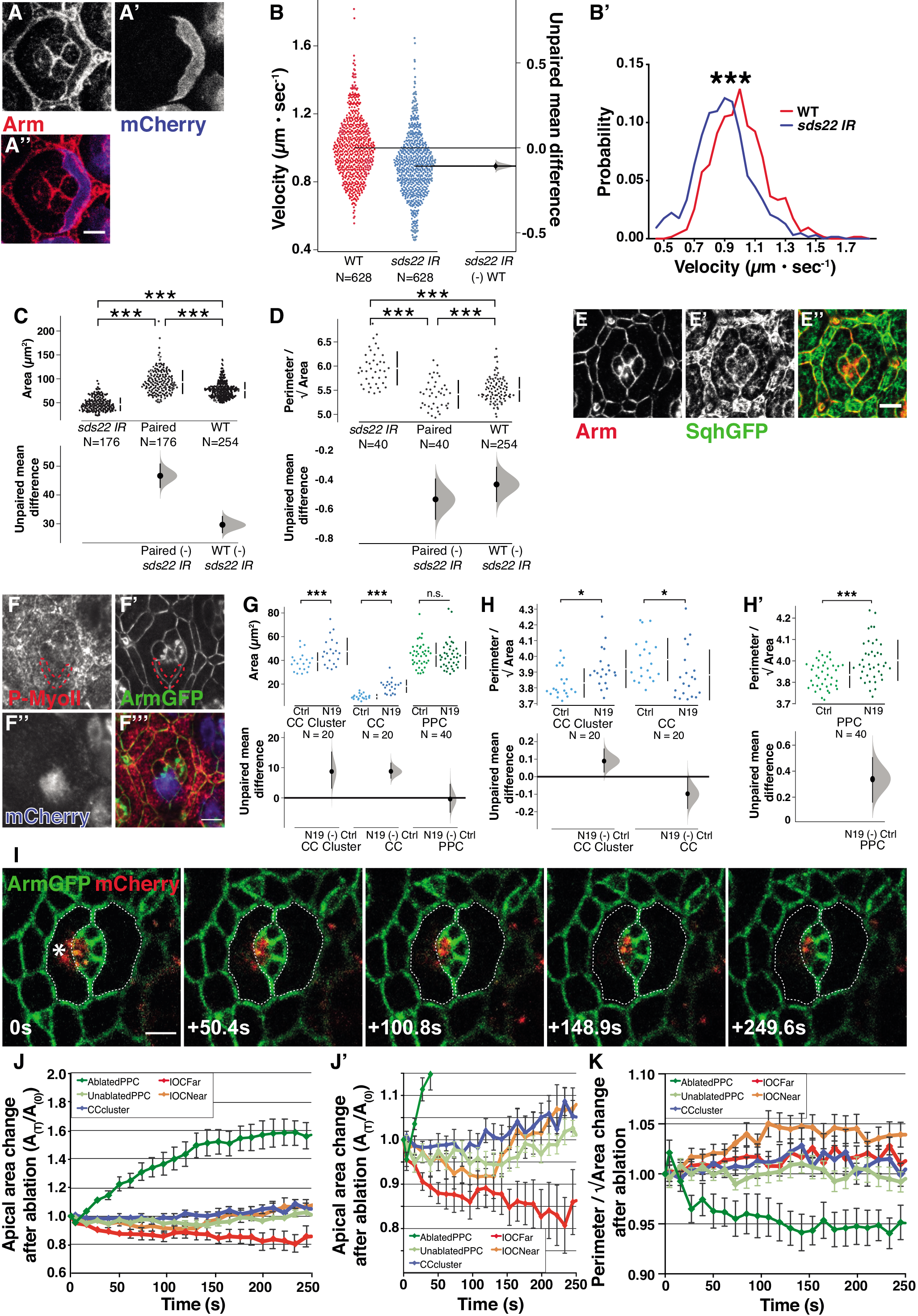
Myosin regulation and mechanical coupling between retinal cells. **(A)** Single primary pigment cell expressing *sds22^IR^*. Clone marked by the presence of mCherry. **(B)** Gardner-Altman estimation plot with left axis showing distribution of velocities of MyoII of wild type and *sds22^IR^* expressing cells. On the right axis, mean difference is plotted as a bootstrap sampling distribution, dot=mean difference, error bars = 95% confidence interval. Unpaired mean difference between WT (n=628) and *sds22^IR^* (n=628) is −0.108 [95CI −0.128; −0.0877]. **(B’)** Velocity distribution for MyoII in wild type versus *sds22^IR^* expressing primary pigment cells (Kolmogorov-Smirnov test, p<0.0001). **(C)** Cummings estimation plot with upper axis showing distribution of areas of primary pigment cells expressing *sds22^IR^*, wild type primary pigment cell in the same ommatidia (paired) and wild-type primary pigment cell in unaffected ommatidia (n= 176, 176, 254 ommatidia respectively, one-way ANOVA, p<0.0001). On the lower axis, mean differences for comparisons to *sds22^IR^* expressing cells are plotted as bootstrap sampling distributions, dot=mean difference, error bars = 95% confidence interval. Unpaired mean difference of: paired (n=176) minus *sds22^IR^* (n=176): 46.7 [95CI 42.4; 50.9]; wild type (n=176) minus *sds22^IR^* (n=254) 29.7 [95CI 26.7; 32.7]. **(D)** Cummings estimation plot with upper axis showing distribution of cell shape index (perimeter/√ area) of primary pigment cells expressing *sds22^IR^*, wild-type primary pigment cell in the same ommatidia (paired) and wild-type primary pigment cell in unaffected ommatidia (n= 40, 40, 90 ommatidia respectively, one-way ANOVA, p<0.0001, *sds22^IR^* −paired p<0.0001, *sds22^IR^* -wild-type p<0.001, paired-wild-type p=0.1585). On the lower axis, mean differences for comparisons to *sds22^IR^* expressing cells are plotted as bootstrap sampling distributions, dot=mean difference, error bars = 95% confidence interval. Unpaired mean difference of: paired (n=40) minus *sds22^IR^* (n=40): −0.538 [95CI −0.678; −0.396]; PPC (n=40) minus *sds22^IR^* (n=90): −0.436 [95CI −0.557; −0.315]. **(E)** Ommatidium from a *sqh^AX3^;sqh::GFP;* retina stained for Arm (Red) showing actomyosin cable around the perimeter of the cone cell cluster. **(F)** Single cone cell expressing Rho^N19^, marked by the presence of mCherry. **(G-H’)** Cummings plots with upper axis showing distribution of **(G)** apical area and **(H-H’)** cell shape index (perimeter/√area) of **(G, H)** cone cell clusters, individual cone cells and **(G, H’)** primary pigment cells in ommatidia with one cone cell expressing Rho^N19^, compared to wild type control ommatidia in the same retinas. For area, one sample T-test: cone cluster p=0.008, cone cell p<0.0001 and primary pigment cell n.s. p=0.8728. For shape index, one sample T-test: cone cluster p=0.0129, cone cell p=0.0465, primary pigment cell p=0.0003. On the lower axes, mean differences for comparisons of cells in Rho^N19^ expressing ommatidia to controls are plotted as bootstrap sampling distributions, dot=mean difference, error bars = 95% confidence interval. Unpaired mean difference of Rho^N19^ expressing minus control for area: cone cell cluster (n=20): 8.74 [95CI 3.06; 15.3]; individual cone cell (n=20): 8.79 [95CI 6.39; 11.7]; primary pigment cell (n=40) minus Control PPC (n=40): −0.397 [95CI −4.93; 4.64]; and for shape index: cone cell cluster (n=20): 0.0894 [95CI 0.0243; 0.161]; individual cone cell (n=20): −0.0974 [95CI −0.184; −0.000475]; primary pigment cell (n=40) minus Control PPC (n=40): 0.338 [95CI 0.158; 0.508]. **(I)** Time-course showing AJs labelled with Arm::GFP after ablation of the medial meshwork in left primary pigment cell in an ommatidium where one cone cell expresses Rho^N19^ (marked by mCherry, red). Region of ablation marked with a white star. Cell outlines before ablation are superimposed on each image with dotted lines. **(J)** Apical area changes over time after ablation, A(T)/A(0), A=apical area (n=9 ommatidia). **(J’)** Magnified panel showing y axis at higher resolution. **(K)** Cell shape index (perimeter/√area) relative to initial shape parameter over time after ablation (n=9 ommatidia). All scale bars = 5 μm.

Secondly, we aimed to test the prediction that the pattern of force propagation in the ommatidium can be explained if the core cone cells are stiffer than the other cell types. Compatible with this idea, the cone cells contain a cortical actomyosin cable that delineates their AJ with the surrounding primary pigment cells (Figure 6E). This could increase cortical tension, and thus make their perimeter stiffer. In order to test this idea, we sought to inhibit actomyosin in the cone cells and assess the impact of this perturbation on force distribution between each lens cell of the ommatidium. To inhibit actomyosin, we expressed dominant negative RhoA (Rho^N19^) in cone cells. This led to an expansion of the cell apical area (Figure 6F-G) and to a distortion of their geometry (Figure 6F, 6H). This shows that actomyosin is required to shape the cone cells. Consistent with our computational model that cone cell stiffness is a parameter that influences the shape of their neighbour, we found that expression of Rho^N19^ in a cone cell induced an elongation of the flanking PPCs (Figure 6H’), without affecting the area of the cell (Figure 6G).

Thirdly, to assess how inhibiting actomyosin contractility in the cone cells affects mechanical coupling between the remaining cells, we performed laser ablation experiments. Ablating the MyoII meshworks in one primary pigment cell in ommatidia in which cone cells express Rho^N19^ did not affect the area and geometry of the cone cell cluster (Figure 6I-K and Movie S8). Quantification of the paired pigment cell revealed that it was not significantly changed, both in area (Figure 6J-J’ and Supplementary Figure 4) and when considering its shape index (Figure 6K). Thus, as predicted by our model, altering the mechanical property of cone cells to render them softer perturbs mechanical coupling between the PPC.

## DISCUSSION

How different cells work together to generate a complex tissue organization is not well understood. Here, we studied tissue organisation in the fly retina, which is made up of different epithelial cell types that organize with crystal-like precision during lens morphogenesis. Our work reveals a preeminent role for the contractile medial MyoII meshworks in regulating the apical area and geometry of the retinal cells. In addition, our results indicate that these contractile meshworks are required for mechanical coupling and communication between cells.

Mechanical coupling between cells has been studied in simple epithelia, where cell packing tends toward regular hexagonal arrays (Gibson *et al.*, 2006; Farhadifar *et al.*, 2007). In these simple tissues, cells maintain similar mechanical properties. Our work shows this is not the case in a more complex epithelia, where cells need to acquire different morphologies. In the eye, our computational model and experimental observations indicate that different cell types have different mechanical properties. The central cone cells do not deform in response to extrinsic forces. Yet by being stiffer than the surrounding cells, they play a role in distributing forces to the neighbouring primary pigment cells. Both our computational model and genetic manipulations suggest this is because the cone cells are stiffer than their neighbours.

In most instances of cell shape changes studied so far, medial MyoII meshworks have been shown to be polarized. In the fly mesoderm, apical constriction is powered by radially polarized contractile meshworks (Martin *et al.*, 2009a). In the germband, pulsatile flows of contractile actomyosin that are polarized along the anterior-posterior axis drive AJ remodelling to promote cell intercalation (Rauzi *et al.*, 2010). In these tissues, where cells are relatively small and present limited numbers of discrete MyoII structures, the calculated time lag between peak contractility and cell apical area fluctuation has been reported to be ~15 secs (Fernandez-Gonzalez and Zallen, 2011; Collinet *et al.*, 2015). In retinal cells, these contractile MyoII meshworks do not appear to be polarized. They exhibit a continuous, distributed, fluctuating mesh through time and space, with multiple contraction nodes that are asynchronous. In these cells, we estimate the time lag between peak contractility and cell apical area fluctuation to be ~30 secs. Further, in contrast to the germband, where intercalating cells maintain their area (Fernandez-Gonzalez and Zallen, 2011; Sawyer *et al.*, 2011) or shrink as they constrict in the mesoderm (Martin *et al.*, 2009a), we find that retinal cells gradually increase their area over time. Therefore, these contractile meshworks can specifically regulate the geometry of apical-junctional cell profile while allowing for area increase.

We speculate that other mechanisms are at play that oppose MyoII contractility. These could include osmotic pressure for example, or preferential adhesion between cells, as demonstrated between the primary pigment and interommatidial cells (Bao *et al.*, 2010; Larson *et al.*, 2010). Finally, we show that in a complex tissue, mechanisms exist that enable cell types to be more resilient to extrinsic forces. This is the case for the cone cells, which show little deformation when challenged with extrinsic mechanical perturbation. Therefore, mechanisms must exist that limit deformation in these cells. One possibility suggested by our model and genetic experiments, is that this can be achieved through increased stiffness.

## MATERIAL AND METHODS

### Fly strains

Flies were raised on standard food at 25°C. Crosses were performed at 25°C. The following fly strains were used:

*; Ecad::GFP;* (Huang *et al.*, 2009).

*sqh^AX3^; sqh>sqhGFP;* (BL #57144, (Royou *et al.*, 2002)).

;Ecad::Tomato; (Huang *et al.*, 2009)).

*; hsflp;;act>CD2>Gal4,UAS-RFP;* (BL #30558).

*; UAS-sds22-RNAi;* (VDRC, 11788).

*; UAS-Rho^N19^;* (BL #58818).

*;; arm - Arm::GFP* (BL #8555). (Orsulic and Peifer, 1996)

To generate single cells deficient for *sds22*, *hs-flp;;actin>CD2>gal4,UAS-RFP* was crossed to *UAS-sds22-RNAi*. Flies were heat shocked at third instar larval stage at 37°C for 10-15min and dissected 4 days later. To generate ommatidia expressing Rho^N19^, *hs-flp;;actin>CD2>gal4,UAS-RFP* was crossed to *UAS-Rho^N19^*. Flies were heat shocked at third instar larval stage at 37°C for 10-15min and staged for laser ablation 2 days later.

### Time-lapse imaging

*;Ecad::GFP;* flies were staged and examined at 15hrs APF at 25°C and the pupal case removed to expose the retina. Pupae were mounted on blue-tac with the retina facing upwards and covered with a coverslip as described in (Fichelson *et al.*, 2012). Timelapse imaging was performed on a Zeiss inverted microscope with an Andor spinning disc using a Plan Neofluar 100x/1.3 Ph3 oil immersion objective. Images were acquired using ImageJ Micromanager software (Edelstein et al., 2014). Retinas were imaged for a minimum of 12hrs acquiring a Z-series in 1μm sections every 5m. Drift in XY and Z was corrected manually. Images were post processed in FIJI using the Stack-reg plugin (Thevenaz *et al.*, 1998) to further correct for drift. For imaging of medial meshwork dynamics, *sqh^AX3^/Y;sqhGFP/EcadTomato;* flies were staged to 20hrs APF, 25hrs APF or 30hrs APF at 25°C and mounted as described above. Retinas were imaged on a Zeiss LSM880 microscope with a Plan Apochromat 63x/NA1.4 oil objective using airyscan detectors. Images were acquired at a 40-45nm pixel size with a speed of 4.35s/frame. Airyscan processing was performed with the Zen software package to increase resolution.

### Measurements of apical area over time

For measurements of the total apical area for each cell type, the outside perimeter of the cone cell cluster, primary pigment cells and interommatidial cells were traced manually using the Freehand selection tool in FIJI on every 20th frame (100mins) of the time-lapse of*; Ecad::GFP;* retinas. Areas enclosed by each perimeter were measured in FIJI and subtracted from each other to generate areas for each cell layer (e.g. area of interommatidial cell outline minus area of primary pigment cell outline gives the area for the interommatidial cell layer). These areas were then expressed as a percentage of the total ommatidium area. 13 ommatidia from the center of the field of view of two independent retinas were registered in time by aligning the mid point of the four-way vertex stage of the cone cell T1 transition and measurements were averaged for each time point.

### Particle Image Velocimetry (PIV)

Time lapse movies of *sqh^AX3^/Y;sqhGFP/EcadTomato;* ommatidia were processed in Fiji with Bleach correction and Gaussian blur, and registered with the ‘Stack-reg’ plugin (Thevenaz *et al.*, 1998) to correct for lateral drift. PIV analysis was performed using the FIJI PIV plugin (Tseng *et al.*, 2012), by choosing an 8×8 pixel window with a time lag 4.34s. Cell contours were tracked using the *Tissue Analyzer* plugin (Aigouy and Le Bivic, 2016) using FIJI, or manually in FIJI to segment the primary pigment, interommatidial and cone cells to measure cell apical areas. For calculation of the advection velocities of medial MyoII, the norms of the PIV vectors were averaged within each cell for each frame and divided by the frame rate in MATLAB. Advection speeds of each cell were then averaged in time for the ommatidia at 20hrs APF, 25hrs APF and 30hrs APF. Mean divergence of velocity vectors within each cell was calculated in MATLAB for each frame. Mean divergence, apical area and apical perimeter measurements over time, were processed by Gaussian smoothing with a window of 43s to improve the signal-to-noise ratio in MATLAB. Cycle lengths and peak amplitude were calculated in MATLAB using the ‘findpeaks’ command. Cross-correlation of the rate of area change and the rate of mean divergence change was performed in MATLAB for each cell and plotted as a heatmap.

### Kymographs

Kymographs were generated using the Reslice plugin in FIJI along a 1 one pixel wide segmented line that was drawn through the center of the cell and did not overlap with the AJ-associated MyoII in any frame. The movie presented in Figure 3 is of a *sqh^AX3^; sqhGFP;* retina at the ‘Late’ stage of development imaged with a Zeiss LSM880 confocal with a Plan Apochromat 63x/NA1.4 oil objective using airyscan detectors drift corrected using the Stackreg plugin (Thévenaz et al., 1998) in FIJI.

### Node density calculation

Sqh::GFP intensity was thresholded on time lapse movies of *sqh^AX3^/Y;sqh::GFP/Ecad::Tomato;* in FIJI to select the nodes of high intensity. The number of nodes per cell and the percentage of cell area covered by MyoII nodes was measured using the ‘Analyze particles’ tool in FIJI and averaged over 50 time-frames for each cell. Multiple retinal cells from 5 independent retinas were analyzed.

### Laser ablation

Ablations of MyoII medial meshworks were performed using a Zeiss LSM880 microscope with a Plan Apochromat 63x/NA1.4 oil objective, using 740nm multiphoton excitation from a Ti-sapphire laser. An ROI of 6×1 pixels (245nm) for 30-40hrs APF retinas and 4×1 pixels (163 nm) for 20hrs APF retinas was drawn in the center of the apical region of the cell and targeted with a laser power of 10-20% at the slowest scan speed for 1 iteration. For monitoring of the MyoII medial meshwork during ablation, *sqh^AX3^/Y;sqh::GFP/Ecad::Tomato;* or *sqh^AX3^/Y;sqh::GFP;* flies were imaged using airyscan detectors. Images were acquired every 3.1-3.6s and processed with the Zen software. For quantification of area change after ablation, *Ecad::GFP* flies were used to visualize *AJs* and images were acquired every 1.27s before and after ablation. Laser ablations in ommatidia expressing dominant negative Rho (Rho^N19^) were performed following the same protocol, except that images were acquired every 2.29s before and after ablation.

### Cell deformation after ablation

Cell deformations after ablation were analyzed on movies of; *Ecad::GFP or sqh^AX3^; SqhGFP* retinas where the primary pigment cell’s medial meshwork was targeted. Each cell type of interest was segmented manually using the Freehand Selection tool in FIJI on the initial frame before ablation and then every 10 frames (12.7s) until the maximum change in area was reached (normally about 200-250s). Area and perimeter/√area were measured and the ratio before and after ablation over time was calculated and averaged across 14 ommatidia. Analysis of cell deformation following laser ablation in ommatidia expressing dominant negative Rho (Rho^N19^) was performed using the same protocol, except that manual segmentation was performed on every 5^th^ frame (11.5s), and measurements were performed on 9 ommatidia.

### Statistical tests

Statistical tests were performed in Graphpad Prism 7. Data were compared using one sample T-test, Kruskal-Wallis test or One-way ANOVA with Tukey’s post-hoc tests. For graphical analysis, data were plot on a Gardner-Altman or Cummings estimation plot using the DABEST package in R with 5000 bootstrap resamples unless otherwise stated. All confidence intervals are bias-corrected and accelerated (Ho *et al.*, 2019)

### Mechanical model for ommatidial morphology

The apical surface of the ommatidial cell cluster is modelled as a composite mechanical medium, consisting of an inner cone cell cluster of area *A_c_*, a medial concentric layer of two adjacent primary pigment cell cells of areas *A*_*p*1_ and *A*_*p*2_, and an outer layer of two interommatidial cell clusters of areas *A*_*i*1_ and *A*_*i*2_ (Figure 5A). In the spirit of vertex models (Farhadifar *et al.*, 2007; Fletcher *et al.*, 2014), each cellular unit (cones, primary or interommatidial) is characterized by a line tension, *γ*, at cell-cell interfaces; a surface tension, Γ, at their interiors; an elastic modulus, *k*, penalizing changes in cell apical area. The edge tension arises from a competition between intercellular adhesion and cortical actomyosin contractility, and the bulk tension arises from contractility in the medial actomyosin meshwork. The total mechanical energy of the cell colony is given by

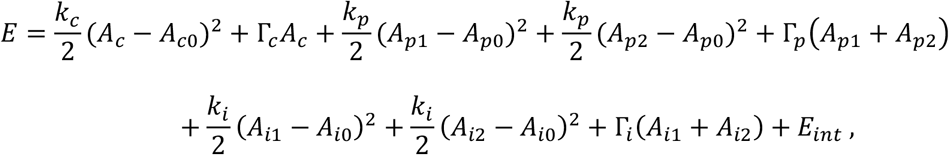

where *k_c_, k_p_*, and *k_i_* define the areal elastic moduli of the cone, primary pigment and interommatidial clusters, Γ_*c*_, Γ_*p*_, and Γ_*i*_ are the respective surface tensions, and *A*_*p*0_, *A*_*i*0_, and *A*_*i*0_ are the respective preferred areas of the three cell types. Intercellular interactions at cell-cell interfaces are given by the term *E_int_*:

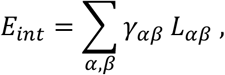

where *γ_αβ_* is the line tension at the interface between cell *α* and *β* (*α,β* ∈ cone cell, primary pigment cell, interommatidial cell) and *L_αβ_* is the length of the interface. Each point, 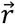, on the cone, primary pigment and interommatidial cell contour evolves in time according to the equation of motion where *μ* is a friction coefficient.

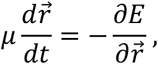

To account for pulsatile actomyosin fluctuations we assume that the cell surface tension fluctuates in time according to, Γ_*α*_(*t*) = Γ_*α*0_ (1 + *θ* sin(*ωt* + *φ_α_*)), where *α* labels the cell type, Γ_*α*0_ is the base tension, *ω* is the frequency of fluctuation, and φ_)_ is a phase-shift calibrated from experimental data. The model is simulated using a custom code implemented using the Surface Evolver program (Brakke, 1992) starting with an initial circular morphology, consisting of an inner circular cone cell cluster, surrounded by concentric rings of primary pigment and interommatidial cells. The model parameters are chosen to reproduce the experimentally measured geometry of the wild type ommatidia (Table 1). To simulate medial meshwork ablation of cell *α*, we let Γ_*α*_ → Γ_*α*_ – ΔΓ, after relaxing the cell system to their mechanical equilibrium state. Similarly, to induce hyper-contractility we let Γ_*α*_ → Γ_*α*_ +ΔΓ after mechanical relaxation.

**Table 1.** Model Parameters.

| Cell type | Preferred area | Surface tension $\Gamma$ | Elastic modulus $k$ |
| --- | --- | --- | --- |
| Primary Pigment (left/right) | 1.75 | 1.0 | 2.5 |
| Cone cell cluster | 1.0 | 1.0 | 14.0 |
| Interommatidial cell cluster (left/right) | 1.5 | 1.0 | 6.0 |
| Interface type | Edge tension $\gamma$ | | |
| cone-primary pigment | 2.5 |  |  |
| primary pigment-primary pigment | 0.5 |  |  |
| primary pigment-interommatidial cells | 2.0 |  |  |
| interommatidial cells-interommatidial cells | 0.5 |  |  |
| interommatidial cells-outside | 2.0 |  |  |

### Immunofluorescence

Whole mount retinas at 40% APF were prepared as previously described (Walther and Pichaud, 2006) The following antibodies were used for indirect immunofluorescence: mouse anti-Arm 1/200 (N27-A1, Developmental Studies Hybridoma Bank) and rabbit anti-P-Myosin Light Chain 2 (S19) 1/50 (3671S, Cell Signalling). Mouse or rabbit secondary antibodies conjugated to Alexa488 or Cy5 (as appropriate) were used at 1/200 each (Jackson ImmunoResearch). Samples were mounted in VectaShield™ and imaging was performed using a Leica SP5 or SP8 confocal microscope. Images were edited using Fiji and Adobe Photoshop 7.0.

## Acknowledgments

We thank the Pichaud lab members, and Yanlan Mao and Melda Tozluoglu for their input during the course of this project. The N2 A71 anti-Armadillo antibody, was deposited to the DSHB by Wieschaus, E. (DSHB Hybridoma Product N2 7A1 Armadillo). Stocks obtained from the Bloomington *Drosophila* Stock Center (NIH P40OD018537) and the Vienna *Drosophila* Resource Center were used in this study. This work was funded by an MRC grant to FP (MC_UU_12018/3) and an MRC PhD studentship to LB. Work in SB lab is funded by Royal Society grant URF\R1\180187, and an EPSRC funded PhD studentship to MFS.

## Author contributions

FP conceived and supervised the work. LB and RFW designed, performed and analyzed the experiments. SB and LB developed analytical tools for experimental data analysis. SB and MFS designed and developed the computational model. FP and LB wrote the manuscript with the help of SB and RFW.

---

**Supplementary Figure 1: Estimating the contraction/relaxation state of retinal cells. (A)** Cummings estimation plot with upper axis showing continuous time spent relaxed or contracted for primary pigment, cone and interommatidial cells (n=9 ommatidia, Kruskal-Wallis test, p<0.0001, post-hoc Dunn’s multiple comparisons tests: primary pigment cells relaxed-primary pigment cell contracted p=0.0007, cone cell relaxed-cone cell contracted p<0.0001, interommatidial relaxed-interommatidial contracted p=0.4585). On the lower axis, mean differences for comparisons to relaxing are plotted as bootstrap sampling distributions, dot=mean difference, error bars = 95% confidence interval. Unpaired mean difference of: PPC contracted (n=80) minus PPC relaxed (n=80): −27.4 [95CI −39.5; −16.1]; CC contracted (n=176) minus CC relaxed (n=168): 26.2 [95CI 17.9; 34.8]; IOC contracted (n=430) minus IOC relaxed (n=427): 5.22 [95CI 0.688; 10]. **(B)** Cumulative mean divergence demonstrating overall contraction or relaxation of each cell type. n=9 ommatidia from 5 pupae, one-sample T-test PPC and CC p>0.0001, IOC p=0.0024.

**Supplementary Figure 2: Intrinsic area fluctuations over short time scales cell responses to ablation.** (**A**) Change in apical area over time after ablation, A(T)/A(0), presented in Figure 4D, but plotted on a log10 scale. A = apical area (n = 14 ommatidia). **(B)** Changes in apical area over time for non-ablated controls, A(T)/A(0), A=apical area. n=9 ommatidia from 5 pupae.

**Supplementary Figure 3: *sds22^IR^* does not affect the AJ pool of MyoII. (A)** SqhGFP intensity in the primary pigment-interommatidial cell AJ for wild type and *sds22^IR^* expressing cells, paired by ommatidia. n=5 ommatidia with one primary pigment cell expressing *sds22 ^IR^* paired to a wild type primary pigment cell, one sample T-test n.s. p=0.846.

**Supplementary Figure 4: Changes in cell area depend upon the actomyosin cytoskeleton. (A)** Change in apical area over time after ablation, A(T)/A(0), upon expression of Rho^N19^, presented in Figure 6J, but plotted on a log10 scale. A = apical area (n = 9 ommatidia).

**Movie S1: Retinal cell apical area change over time.** *;Ecad::GFP*; pupal retina with AJs labelled in grey by Ecad::GFP. Imaging initiated from 15hrs APF for ~12hrs showing how ommatidial cells change shape and size during development. 5min frame interval. Movie post-processed by bleach correction and filtering with a Gaussian blur in FIJI. Scale bar = 5μm

**MovieS2: MyoII dynamics in retinal cells.** Airyscan processed time-lapse of *sqh^AX3^;sqh::GFP/Ecad::Tomato;* pupal retina at 30hrs APF. Sqh::GFP in green and AJs in red marked by Ecad::Tomato. Example of movie used for PIV analysis. 4.35sec frame interval. Scale bar = 5μm

**MovieS3: MyoII dynamics in retinal cells.** Airyscan processed time-lapse of *sqh^AX3^;sqh::GFP/Ecad::Tomato;* pupal retina at 30hrs APF showing example region of the primary pigment cell. Sqh::GFP in green and AJs in red marked by Ecad::Tomato. Note that area constrictions occur in tandem with Sqh::GFP pulses. Scale bar = 5μm

**MovieS4: Medial meshwork recovers after ablation and restores cell area.** Airyscan processed time lapse of *sqh^AX3^;sqh::GFP;* pupal retina with Sqh::GFP in grey, showing MyoII dynamics after ablation of the medial meshwork in the left-hand primary pigment cell in frame 3 and then subsequent repair of the meshwork. Note how the cell relaxes area and loses shape but that this is regained as the meshwork repairs. 3.55sec frame interval. Scale bar = 5μm

**MovieS5: MyoII medial meshwork ablation in primary pigment cell.** Airyscan processed timelapse of *sqh^AX3^;sqh::GFP;* pupal retina with Sqh::GFP in grey, showing MyoII dynamics after ablation of the MyoII medial meshwork in the left-hand primary o o pigment cell. Ablation occurs at frame 3 (3.45sec). 1.15sec frame interval. Scale bar = 5μm

**MovieS6: Cell shape changes after MyoII medial meshwork external force perturbation.** Time-lapse of *;Ecad::GFP;* pupal retina at 30hrs APF with AJs in grey marked with Ecad::GFP. Ablation of medial meshwork in left primary pigment cell occurs at frame 5 (5.08sec). 1.27sec frame interval. Note how neighbouring cell types vary in their response to ablation in terms of area and shape changes. Movie postprocessed by filtering with a Gaussian blur in FIJI. Scale bar = 5μm

**MovieS7: Laser cutting of one of the AJ shared by the two primary pigment cells.** Representative laser ablation of a primary-primary pigment cell AJ in an ommatidium at 20hrs AFP. Cell outlines are labelled with ECad::GFP. Note how junction repairs by end of movie. Frame interval: 0.93sec. Scale bar = 5μm.

**MovieS8: MyoII medial meshwork ablation in primary pigment cell containing a Rho^N19^ cone cells.** Airyscan processed time-lapse of ArmGFP pupal retina. The cone cells expressing Rho^N19^ is labelled using mCherryFP. The ablation occurs at frame 3 (6.87sec). 2.29sec frame interval. Scale bar = 5μm

## REFERENCES

Aigouy, B., and Le Bivic, A. (2016). The PCP pathway regulates Baz planar distribution in epithelial cells. Sci Rep 6, 33420.

Bao, S., Fischbach, K.F., Corbin, V., and Cagan, R.L. (2010). Preferential adhesion maintains separation of ommatidia in the Drosophila eye. Dev Biol 344, 948–956.

Bertet, C., Sulak, L., and Lecuit, T. (2004). Myosin-dependent junction remodelling controls planar cell intercalation and axis elongation. Nature 429, 667–671.

Bi, D., Lopez, J., Schwarz, J. Manning, L. (2015). A density-independent rigidity transition in biological tissues.. Nature Phys 11, 1074–1079.

Blankenship, J.T., Backovic, S.T., Sanny, J.S., Weitz, O., and Zallen, J.A. (2006). Multicellular rosette formation links planar cell polarity to tissue morphogenesis. Dev Cell 11, 459–470.

Brachmann, C.B., and Cagan, R.L. (2003). Patterning the fly eye: the role of apoptosis. Trends Genet 19, 91–96.

Brakke, K.A. (1992). The Surface Evolver. Experimental Mathematics 1, 14.

Chan, E.H., Chavadimane Shivakumar, P., Clement, R., Laugier, E., and Lenne, P.F. (2017). Patterned cortical tension mediated by N-cadherin controls cell geometric order in the Drosophila eye. Elife 6.

Collinet, C., Rauzi, M., Lenne, P.F., and Lecuit, T. (2015). Local and tissue-scale forces drive oriented junction growth during tissue extension. Nat Cell Biol 17, 1247–1258.

Coravos, J.S., Mason, F.M., and Martin, A.C. (2017). Actomyosin Pulsing in Tissue Integrity Maintenance during Morphogenesis. Trends Cell Biol 27, 276–283.

Duda, M., Kirkland, N.J., Khalilgharibi, N., Tozluoglu, M., Yuen, A.C., Carpi, N., Bove, A., Piel, M., Charras, G., Baum, B., and Mao, Y. (2019). Polarization of Myosin II Refines Tissue Material Properties to Buffer Mechanical Stress. Dev Cell 48, 245–260 e247.

Farhadifar, R., Roper, J.C., Aigouy, B., Eaton, S., and Julicher, F. (2007). The influence of cell mechanics, cell-cell interactions, and proliferation on epithelial packing. Curr Biol 17, 2095–2104.

Fernandez-Gonzalez, R., and Zallen, J.A. (2011). Oscillatory behaviors and hierarchical assembly of contractile structures in intercalating cells. Phys Biol 8, 045005.

Fichelson, P., Brigui, A., and Pichaud, F. (2012). Orthodenticle and Kruppel homolog 1 regulate Drosophila photoreceptor maturation. Proc Natl Acad Sci U S A 109, 7893–7898.

Fletcher, A.G., Osterfield, M., Baker, R.E., and Shvartsman, S.Y. (2014). Vertex models of epithelial morphogenesis. Biophys J 106, 2291–2304.

Gibson, M.C., Patel, A.B., Nagpal, R., and Perrimon, N. (2006). The emergence of geometric order in proliferating metazoan epithelia. Nature 442, 1038–1041.

Gomez-Galvez, P., Vicente-Munuera, P., Tagua, A., Forja, C., Castro, A.M., Letran, M., Valencia-Exposito, A., Grima, C., Bermudez-Gallardo, M., Serrano-Perez-Higueras, O., Cavodeassi, F., Sotillos, S., Martin-Bermudo, M.D., Marquez, A., Buceta, J., and Escudero, L.M. (2018). Scutoids are a geometrical solution to threedimensional packing of epithelia. Nat Commun 9, 2960.

Grusche, F.A., Hidalgo, C., Fletcher, G., Sung, H.H., Sahai, E., and Thompson, B.J. (2009). Sds22, a PP1 phosphatase regulatory subunit, regulates epithelial cell polarity and shape [Sds22 in epithelial morphology]. BMC Dev Biol 9, 14.

Hayashi, T., and Carthew, R.W. (2004). Surface mechanics mediate pattern formation in the developing retina. Nature 431, 647–652.

Heisenberg, C.P., and Bellaiche, Y. (2013). Forces in tissue morphogenesis and patterning. Cell 153, 948–962.

Ho, J., Tumkaya, T., Aryal, S., Choi, H., and Claridge-Chang, A. (2019). Moving beyond P values: data analysis with estimation graphics. Nature methods 16, 565–566.

Huang, J., Zhou, W., Dong, W., Watson, A.M., and Hong, Y. (2009). From the Cover: Directed, efficient, and versatile modifications of the Drosophila genome by genomic engineering. Proc Natl Acad Sci U S A 106, 8284–8289.

Kafer, J., Hayashi, T., Maree, A.F., Carthew, R.W., and Graner, F. (2007). Cell adhesion and cortex contractility determine cell patterning in the Drosophila retina. Proc Natl Acad Sci U S A 104, 18549–18554.

Kasza, K.E., Farrell, D.L., and Zallen, J.A. (2014). Spatiotemporal control of epithelial remodeling by regulated myosin phosphorylation. Proc Natl Acad Sci U S A 111, 11732–11737.

Larson, D.E., Johnson, R.I., Swat, M., Cordero, J.B., Glazier, J.A., and Cagan, R.L. (2010). Computer simulation of cellular patterning within the Drosophila pupal eye. PLoS computational biology 6, e1000841.

Lavoie, J., Gasso Astorga, P., Segal-Gavish, H., Wu, Y.C., Chung, Y., Cascella, N.G., Sawa, A., and Ishizuka, K. (2017). The Olfactory Neural Epithelium As a Tool in Neuroscience. Trends in molecular medicine 23, 100–103.

Lecuit, T., and Yap, A.S. (2015). E-cadherin junctions as active mechanical integrators in tissue dynamics. Nat Cell Biol 17, 533–539.

Martin, A.C., Kaschube, M., and Wieschaus, E.F. (2009a). Pulsed contractions of an actin-myosin network drive apical constriction. Nature 457, 495–499.

Martin, R., Smibert, P., Yalcin, A., Tyler, D.M., Schafer, U., Tuschl, T., and Lai, E.C. (2009b). A Drosophila pasha mutant distinguishes the canonical microRNA and mirtron pathways. Mol Cell Biol 29, 861–870.

Mason, F.M., Xie, S., Vasquez, C.G., Tworoger, M., and Martin, A.C. (2016). RhoA GTPase inhibition organizes contraction during epithelial morphogenesis. J Cell Biol 214, 603–617.

Munjal, A., and Lecuit, T. (2014). Actomyosin networks and tissue morphogenesis. Development 141, 1789–1793.

Munjal, A., Philippe, J.M., Munro, E., and Lecuit, T. (2015). A self-organized biomechanical network drives shape changes during tissue morphogenesis. Nature 524, 351–355.

Orsulic, S., and Peifer, M. (1996). An in vivo structure-function study of armadillo, the beta-catenin homologue, reveals both separate and overlapping regions of the protein required for cell adhesion and for wingless signaling. J Cell Biol 134, 1283–1300.

Rauzi, M., Krzic, U., Saunders, T.E., Krajnc, M., Ziherl, P., Hufnagel, L., and Leptin, M. (2015). Embryo-scale tissue mechanics during Drosophila gastrulation movements. Nat Commun 6, 8677.

Rauzi, M., Lenne, P.F., and Lecuit, T. (2010). Planar polarized actomyosin contractile flows control epithelial junction remodelling. Nature 468, 1110–1114.

Ready, D.F. (1989). A multifaceted approach to neural development. Trends Neurosci 12, 102–110.

Roh-Johnson, M., Shemer, G., Higgins, C.D., McClellan, J.H., Werts, A.D., Tulu, U.S., Gao, L., Betzig, E., Kiehart, D.P., and Goldstein, B. (2012). Triggering a cell shape change by exploiting preexisting actomyosin contractions. Science 335, 1232–1235.

Royou, A., Sullivan, W., and Karess, R. (2002). Cortical recruitment of nonmuscle myosin II in early syncytial Drosophila embryos: its role in nuclear axial expansion and its regulation by Cdc2 activity. J Cell Biol 158, 127–137.

Rupprecht, J.F., Ong, K.H., Yin, J., Huang, A., Dinh, H.H., Singh, A.P., Zhang, S., Yu, W., and Saunders, T.E. (2017). Geometric constraints alter cell arrangements within curved epithelial tissues. Mol Biol Cell 28, 3582–3594.

Sawyer, J.K., Choi, W., Jung, K.C., He, L., Harris, N.J., and Peifer, M. (2011). A contractile actomyosin network linked to adherens junctions by Canoe/afadin helps drive convergent extension. Mol Biol Cell 22, 2491–2508.

Thevenaz, P., Ruttimann, U.E., and Unser, M. (1998). A pyramid approach to subpixel registration based on intensity. IEEE Trans Image Process 7, 27–41.

Tseng, Q., Duchemin-Pelletier, E., Deshiere, A., Balland, M., Guillou, H., Filhol, O., and Thery, M. (2012). Spatial organization of the extracellular matrix regulates cellcell junction positioning. Proc Natl Acad Sci U S A 109, 1506–1511.

Turlier, H., and Betz, T. (2019). Unveiling the Active Nature of Living-Membrane Fluctuations and Mechanics. Annual Review of Condensed Matter Physics 10, 213–232.

Vasquez, C.G., Tworoger, M., and Martin, A.C. (2014). Dynamic myosin phosphorylation regulates contractile pulses and tissue integrity during epithelial morphogenesis. J Cell Biol 206, 435–450.

Walther, R.F., and Pichaud, F. (2006). Immunofluorescent staining and imaging of the pupal and adult Drosophila visual system. Nat Protoc 1, 2635–2642.

