## Supplementary figures and images for "Cell-type specific mechanical response and myosin dynamics during retinal lens development in *Drosophila*"

### Figure S1

**Supplementary Figure 1: Estimating the contraction/relaxation state of retinal cells.**

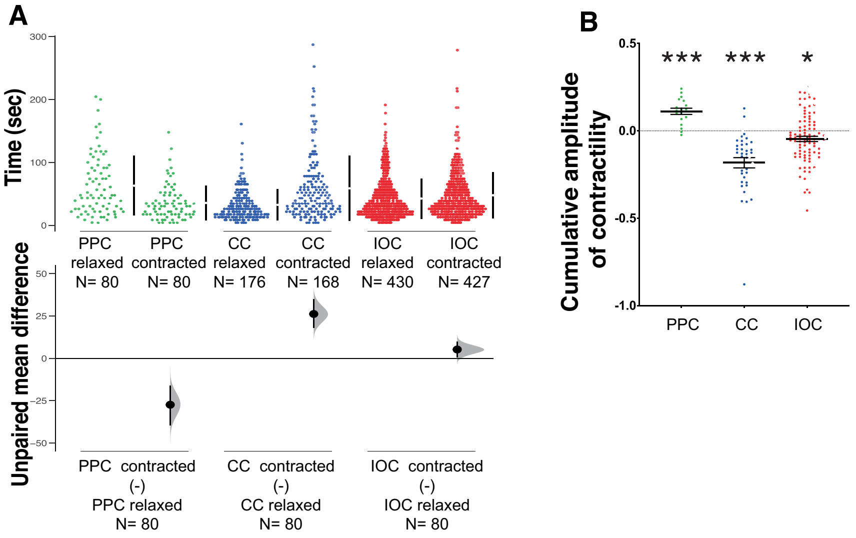

### Figure S2

#### Supplementary Figure 2: Intrinsic area fluctuations and cell responses to ablation

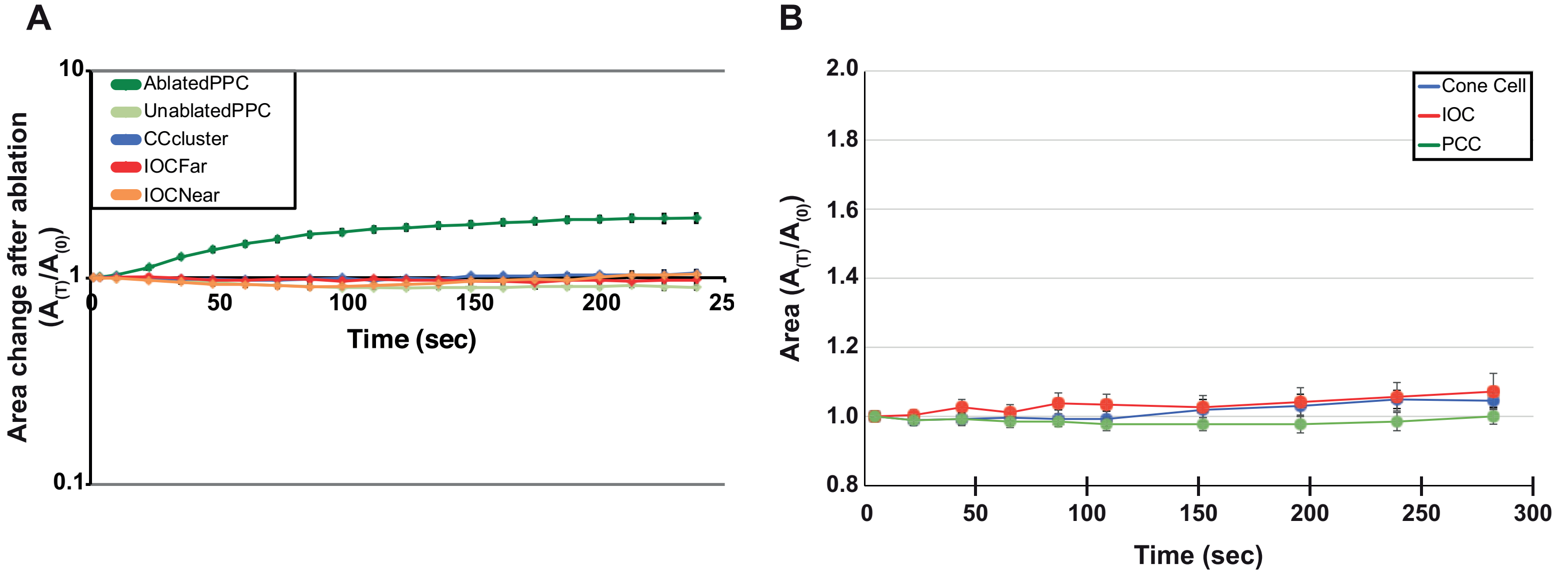

### Figure S3

**Supplementary Figure 3: *sds22$^{IR}$* does not affect the AJ pool of MyoII**

**A**

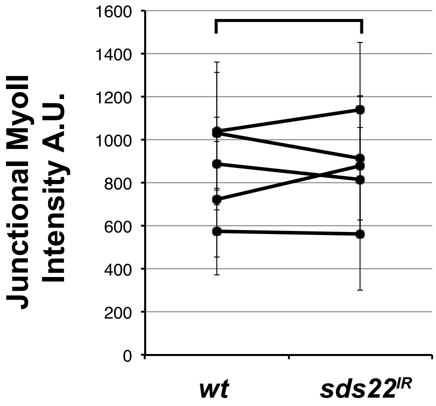

### Figure S4

**Supplementary Figure 4: Changes in cell area depend upon the actomyosin cytoskeleton**

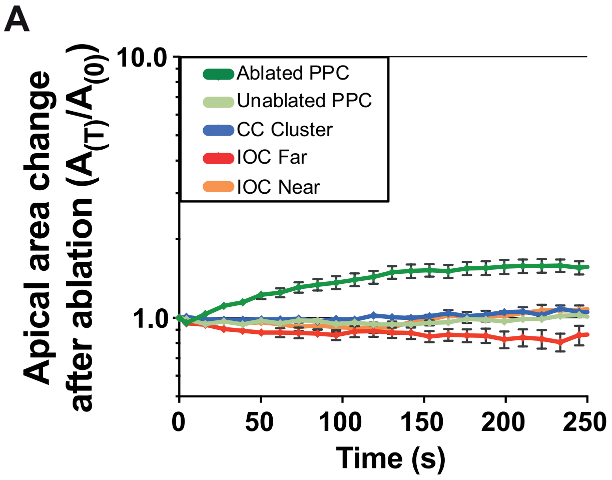
